# Discrimination of regular and irregular rhythms by accumulation of time differences

**DOI:** 10.1101/2020.07.04.187708

**Authors:** Marisol Espinoza-Monroy, Victor de Lafuente

## Abstract

Perceiving the temporal regularity in a sequence of repetitive sensory events facilitates the preparation and execution of relevant behaviors with tight temporal constraints. How we estimate temporal regularity from repeating patterns of sensory stimuli is not completely understood. We developed a decision-making task in which participants had to decide whether a train of visual, auditory, or tactile pulses, had a regular or an irregular temporal pattern. We tested the hypothesis that subjects categorize stimuli as irregular by accumulating the time differences between the predicted and observed times of sensory pulses defining a temporal rhythm. Results show that instead of waiting for a single large temporal deviation, participants accumulate timing-error signals and judge a pattern as irregular when the amount of evidence reaches a decision threshold. Model fits of bounded integration showed that this accumulation occurs with negligible leak of evidence. Consistent with previous findings, we show that participants perform better when evaluating the regularity of auditory pulses, as compared with visual or tactile stimuli. Our results suggest that temporal regularity is estimated by comparing expected and measured pulse onset times, and that each prediction error is accumulated towards a threshold to generate a behavioral choice.

## Introduction

A sequence of repeating sensory pulses is often perceived as a rhythm, allowing the observer to predict the onset of future stimuli and synchronize relevant motor plans accordingly (Cadena-Valencia et al., 2018; Gámez et al., 2018; Itoh et al., 2020; Remington et al., 2018). How the brain perceives rhythm from a train of repeating sensory pulses is not completely understood (Finnerty et al., 2015; Ivry, 1996; Lakatos et al., 2019; Mauk & Buonomano, 2004; Merchant et al., 2013; Paton & Buonomano, 2018). Previous work has shown that regularly repeating stimuli gives rise to expectation neuronal activity that encodes the estimated time of future stimuli (Cadena-Valencia et al., 2018; Egger et al., 2019; Gámez et al., 2019; Leon & Shadlen, 2003; Patel & Iversen, 2014; Renoult et al., 2006; Rossi-Pool et al., 2019; Takeya et al., 2017; Toiviainen et al., 2020). The ability to predict the time of future stimuli allows the brain furnishing error signals that quantify the difference between predicted and actual time of stimulus onset (Egger et al., 2019; Ohmae & Tanaka, 2016; Rhodes & Di Luca, 2016). We reasoned that subjects might be able to use the difference between expected and actual onset times to decide whether a train of sensory pulses has a regular or an irregular temporal structure. Under this hypothesis, an observer perceives a stimulus as irregular when the difference between predicted and actual onset times consistently gives evidence in favor of irregular temporal intervals between sensory pulses.

The bounded accumulation of evidence is a mechanism that successfully explains decision-making under varied sensory contexts such as the perception of visual motion, and the discrimination of auditory and tactile stimuli (Brunton et al., 2013; Churchland et al., 2008; de Lafuente et al., 2015; Hernández-Pérez et al., 2020; Pinto et al., 2018; Ratcliff et al., 2016). However, it is not known whether a sequence of consecutive temporal errors can be accumulated by an observer to discriminate between regular and irregular temporal structures. Under the framework of timing-error accumulation there are two extreme modes of operation to generate a decision: (1) subjects wait for a large temporal deviation to judge a rhythm as irregular (Stine et al., 2020; Waskom & Kiani, 2018); or, conversely (2) they accumulate each temporal deviation, and make a decision when the summed evidence reaches a decision threshold.

We aimed to test whether bounded accumulation provided an adequate description of the decision process about temporal regularity, and we also determined which of the two decision strategies participants use to generate perceptual judgements. We tested the generality of this decision model by making use of visual, auditory, and tactile trains of stimuli. Our results show that human participants are able to accumulate temporal prediction errors and judge a train of sensory pulses as irregular when the accumulated deviations reach a decision bound. Conversely, a rhythm is judged regular by the absence of consistent timing errors, and the decision time is determined by an urgency signal modulating the decision speed-accuracy trade off.

## Methods

### Regularity discrimination task

To characterize the ability of human subjects to distinguish regular from irregular temporal patterns we asked them to perform a discrimination task in which they had to perceive trains of brief (50 ms) sensory pulses and decide whether they appeared at irregular or irregular intervals (Figure 1a) (Buhusi et al., 2016; Celma-Miralles & Toro, 2020; Schulze, 1989). The variability of the interpulse intervals was the key experimental variable determining task difficulty. In this manner, intervals with zero standard deviation defined *regular* stimuli, and any non-zero variability defined *irregular* stimuli. Across trials, the standard deviation of interval durations ranged from 0% (completely regular stimuli) up to 22.9% (highly irregular stimuli) with respect to the mean interval (Figure 1b). We used three mean intervals in order to cover a wide temporal scale of the stimuli (0.35, 0.65, and 1.25 s; randomly selected). Importantly, participants were free to indicate their decision (*regular* or *irregular*) at any time during the presentation of the stimuli, thus defining a *reaction-time discrimination task*. Performance in the discrimination task was characterized by the *psychometric* curve (Kingdom & Prins, 2016), which depicts the proportion of *irregular* responses as a function of irregularity (standard deviation) expressed in time units (seconds), or expressed in percentage with respect to mean interval (Figure 2a,c,e). *Chronometric* curves were used to show decision response time as a function of irregularity (Figure 2b,d,e).

**Figure 1.**
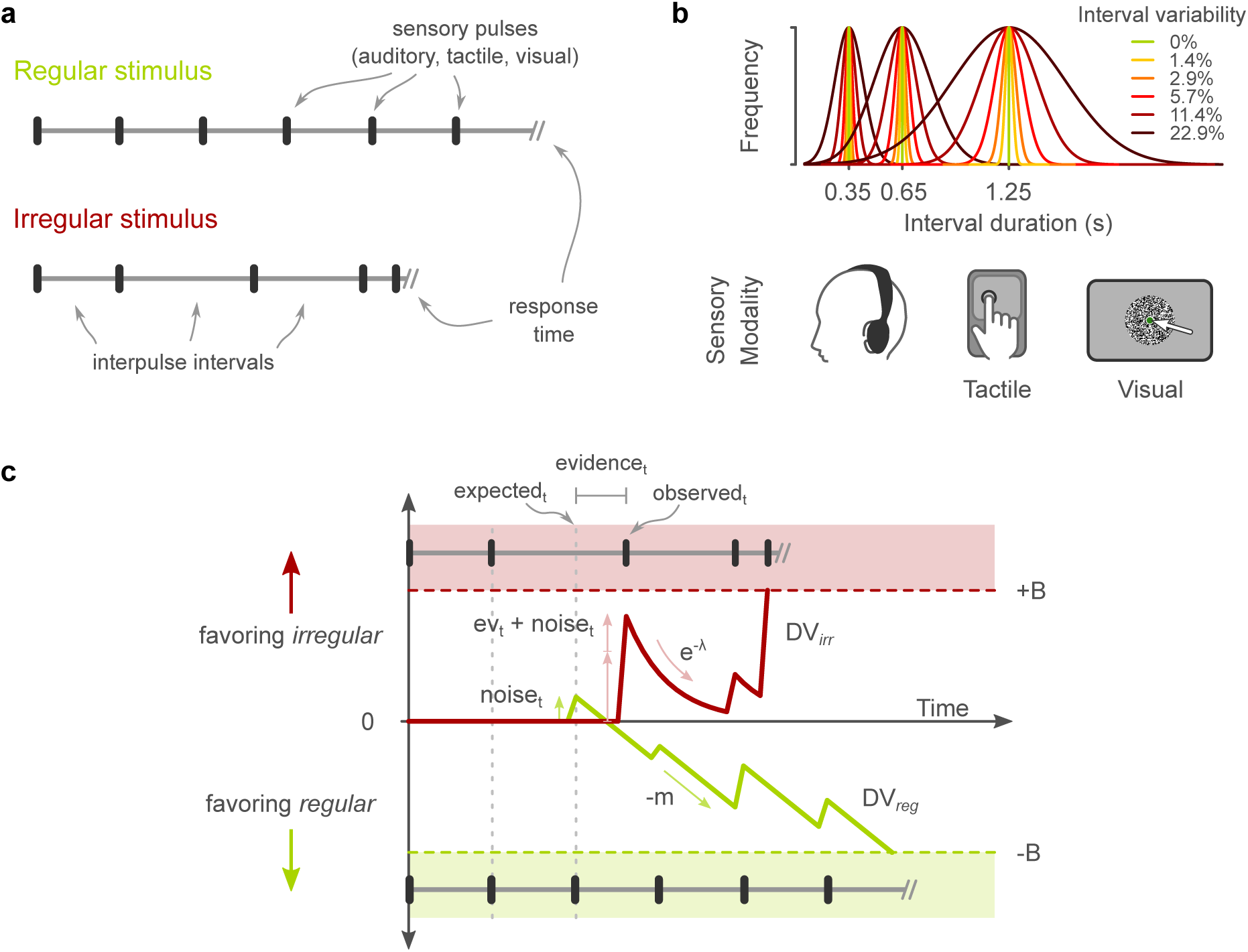
Regularity discrimination task and model of decision making. **(a)** Participants sensed trains of sensory pulses to decide whether they had regular or irregular interpulse intervals. A regular and an irregular pulse train is shown. The sensory pulses consisted of 50-ms auditory, visual, or tactile stimuli. Response times were freely determined by the participants. **(b)** One of three mean interval durations was pseudo-randomly chosen on each trial (0.35, 0.65, 1.25 s). Within a trial, successive intervals were either constant (*regular* trials; 0% variability) or chosen from a distribution with 1.4, 2.9, 5.7, 11.4, or 22.9% standard deviation of the mean (i*rregular* trials). **(c)** Model of decision-making. Two competing decision variables represent evidence in favor of irregular (*DV*_*irr*_) or regular (*DV*_*reg*_) stimuli. The first decision variable to reach its threshold determines the trial’s choice and response time. The irregularity evidence plus a noise term move *DV*_*irr*_ towards the upper bound (+B), favoring an irregular decision. An exponential decay parameter determines how fast the accumulated evidence decreases before the arrival of the next sensory pulse (*λ*). The competing process *DV*_*reg*_ moves towards the lower bound (-B) at a speed determined by -*m*, broken by upward jumps from a noise term (see Equations 2 and 3).

**Figure 2.**
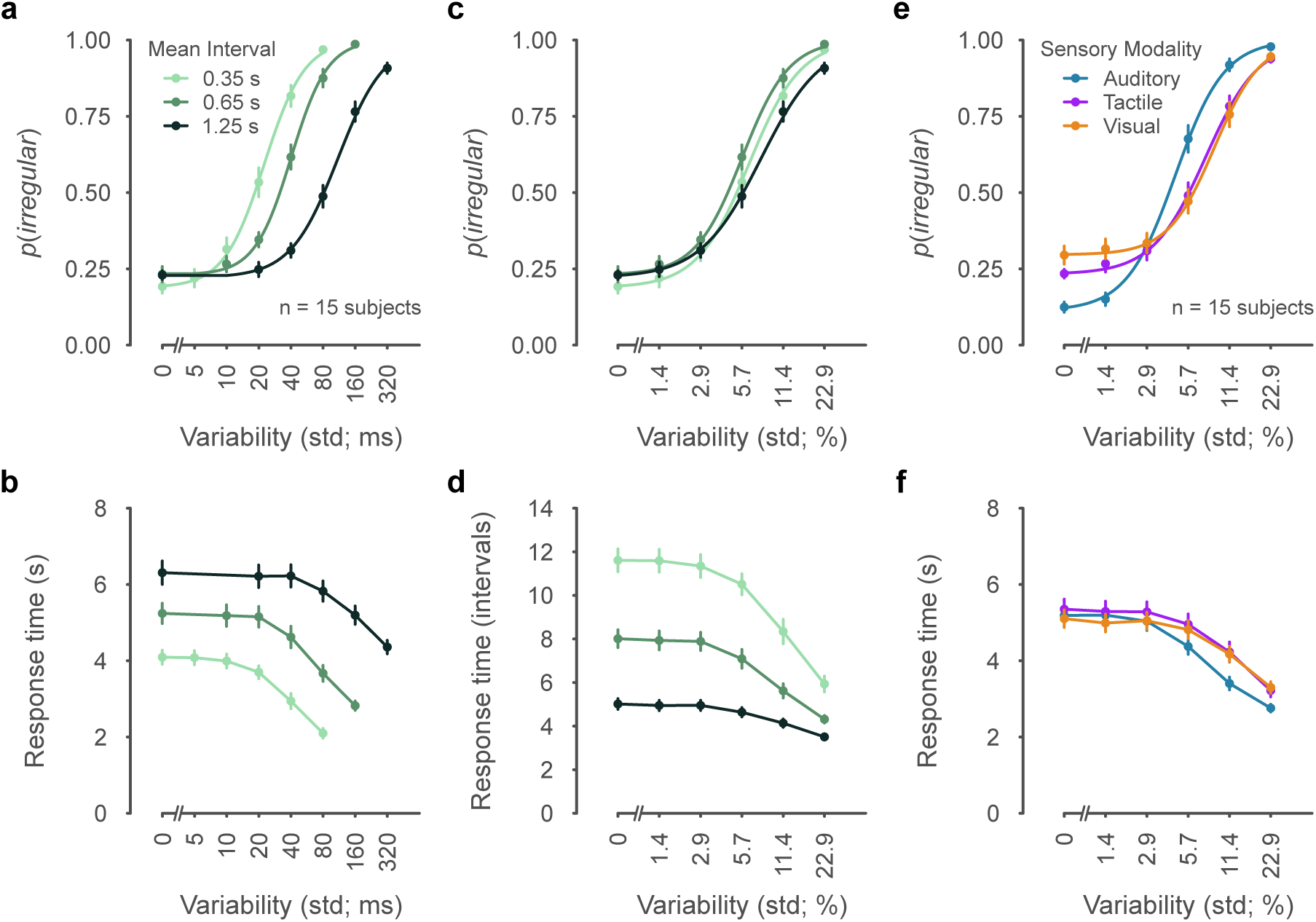
Behavioral choices and response times on the discrimination task. **(a)** Psychometric curves depict the probability of an *irregular* decision as a function of interval variability (standard deviation of interval duration, std, in ms units). Data is conditioned by mean interval (color coded). Dots and error bars denote mean ±s.e.m (n=15 participants). **(b)** Chronometric curves show response times as a function of interval variability (ms units). **(c)** Psychometric curves with variability expressed as percentage of interval duration (% units). **(d)** Chronometric curves in which response times are expressed as number of elapsed intervals. **(e)** Psychometric curves conditioned by sensory modality (color coded). **(f)** Chronometric curves by sensory modality. Response times expressed in seconds and variability in % units.

Trials initiated with a fixation point (2°) at the center of the screen, its color indicating the sensory modality of the upcoming sensory pulses (blue, green, or pink indicated auditory, visual, or tactile pulses, respectively; Figure 1b). The train of sensory pulses started after a random delay (2-3 s) after the fixation point. Participants were instructed to attend to the temporal pattern of the pulses and decide whether they appeared at regular or irregular intervals. The train of sensory pulses stopped after the participants pressed one of the two keys to indicate their decision (F and S keys to indicate a *regular* or an *irregular* choice, respectively). Feedback was provided by presenting the words “correct” or “incorrect” on the screen. A new trial started after a random inter-trial delay (3-4 s). The mean interpulse interval was pseudo-randomly chosen on each trial (0.35, 0.65, or 1.25 s). The standard deviation of the interpulse interval, a term that we name *irregularity*, was pseudo-randomly chosen from the values 0, 1.4, 2.9, 5.7, 11.4 and 22.9% (relative to the mean interval). To obtain a balanced prior probability of *regular* and *irregular* decisions, half the trials had 0% irregularity, and the other half had one of the non-zero irregularities. The temporal pattern presented on each trial was thus constructed by pseudo-randomly selecting (1) the sensory modality, (2) the mean, (3) and the standard deviation of the interpulse intervals, with the restriction that intervals should not be shorter than 50 ms. Trials with a response time larger than 20 intervals, or shorter than 2 intervals, were repeated later in the session without the subject’s knowledge.

Auditory pulses were presented through headphones and consisted of a 50-ms 0.5-kHz sinusoidal tone. Visual pulses were defined by a circle of pixel noise (8.2°), presented for 50 ms behind the fixation point (3 frames of dynamic pixel noise). Tactile pulses were presented through a custom-made electro-magnet stimulator driven by amplifying the voltage of computer’s audio output. Participants placed their right index finger over the rounded stimulator probe (3 mm in diameter) which protruded 3 mm over the rest surface (Figure 1b). Participants wore headphones delivering masking white noise on every trial. Fifteen human subjects (age range: 22-32; 6 female) participated in the study. Each participant completed 15-18 sessions (90 trials each, 15 to 20 min per session) over the course of 5-6 days. They were informed about the purpose and methods of the experiment and provided their written consent. The experimental protocol was approved by the Ethics in Research Committee of the Institute of Neurobiology of National Autonomous University of Mexico. Stimuli presentation and data collection were performed with a computer setup running Matlab and the Psychophysics Toolbox ((Brainard, 1997); monitor: Dell SE2717H, Full HD 1980×1080, 60 Hz; computer: Dell Precision Tower 5810, Windows 10 Pro, 64 bits, NVIDIA Quadro K420 2GB graphics card).

### Decision-making model of temporal pattern discrimination

To test the hypothesis that subjects integrate timing irregularities we used the *accumulation-to-bound* framework of decision-making. Under this framework, an *irregular* decision is made when an *irregular* decision variable (*DV*_*irr*_) reaches the upper decision bound (Figure 1c). *Regular* decisions were modeled by a competing decision variable (*DV*_*reg*_) that moved towards the lower bound as a function of elapsed time (Figure 1c). In this *two-race model* the behavioral decision (*regular* or *irregular*) is determined by the first process to reach its decision bound (Figure 1c). A train of highly irregular pulses would provide evidence rapidly moving *DV*_*irr*_ upwards to the irregular bound. Conversely, a train of regular pulses would fail to provide irregularity evidence, allowing *DV*_*reg*_ to reach the lower decision bound.

An alternative hypothesis to the accumulation model (often termed *independent sampling*) proposes that no accumulation takes place, and decisions are driven by sequentially comparing the most recent sensory input to the decision bound. Under this alternative and *irregular* decision would be made by a given interval deviating more than a defined threshold. The extent to which *accumulation* or *independent sampling* better explains the participants’ behavioral responses can be estimated by a decay (leak) parameter multiplying *DV*_*irr*_ (Figure 1c). If the decay of irregularity evidence is slow with respect to interval duration, no evidence is lost between pulses and irregularity evidence is accumulated perfectly. Conversely, a fast decay term would result in irregularity evidence being lost before the arrival of next sensory pulse, preventing accumulation so that only the most recent evidence would be used to make a decision. In our model, *DV*_*irr*_ and *DV*_*reg*_ are updated at each sensory pulse, starting with the arrival of the third pulse. This is because the third sensory pulse defines the second interval, which is the first interval that can be compared with the previous one. The model assumes that subjects calculate irregularity evidence by comparing the duration of each new interval with the duration of the previous one. Irregularity evidence is thus defined by the absolute difference between the intervals, normalized by the previous interval. It is important to note that there is no evidence favoring regularity, i.e., it is the absence of irregularity that allows *DV*_*reg*_ to reach the lower bound to generate a *regular* stimulus choice. If the subscript *t* denotes each interval, irregularity evidence is defined as:

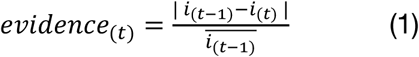

where *i*_(*t*−1)_ − *i*_(*t*)_ is the difference between the duration of the current and previous interval. The evidence provided by each new interval is summed to the previous evidence (along with noise) to define *DV*_irr_ as follows:

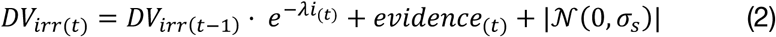

where |𝒩(0, σ_*s*_)| is positive noise parametrized by σ_*s*_; *i*_(*t*)_ is the duration of the new interval, and *λ* is the decay constant of the accumulated evidence. *DV*_*reg*_ is modeled as a downward slope broken by upwards jumps of positive noise as follows:

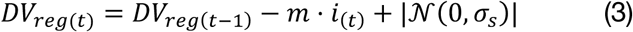

where *m* is the slope parameter and *i*_(*t*)_ is the duration of the new interval (Figure 1c). *Regular* or *irregular* model choices were determined by the first decision variable to reach its bound (+B,-B, respectively) and response time was defined as the number of elapsed intervals after the first sensory pulse.

The four model parameters defining our model (decision bound height *B*, evidence decay constant *λ*, slope *m*, and noise σ_*s*_) were fitted to the psychometric and chronometric curves by means of a grid search within parameter ranges determined by an initial exploratory search (Gherman & Philiastides, 2018). Each parameter combination within the grid search instantiated a model that was run for 500 trials for each percentage of irregularity, thus obtaining the model’s *psychometric* and *chronometric* curves that were compared with the subjects’ corresponding curves. The parameters that minimized the difference between the behavioral and model curves were obtained for each subject and experimental condition. It is important to note that *psychometric* and *chronometric* curves are fit simultaneously, i.e., a 4-tuple of parameters defines both curves. Given that *chronometric* and *psychometric* curves have different y-scales, they were standardized to values between 0 and 1 before calculating the fitting error. To fit the behavioral data of each participant, trials across modalities were pooled to obtain the parameters for the 3 interval durations (Figure 4). Conversely, interval durations were pooled to obtain the parameters for the 3 sensory modalities (Figure 4).

**Figure 3.**
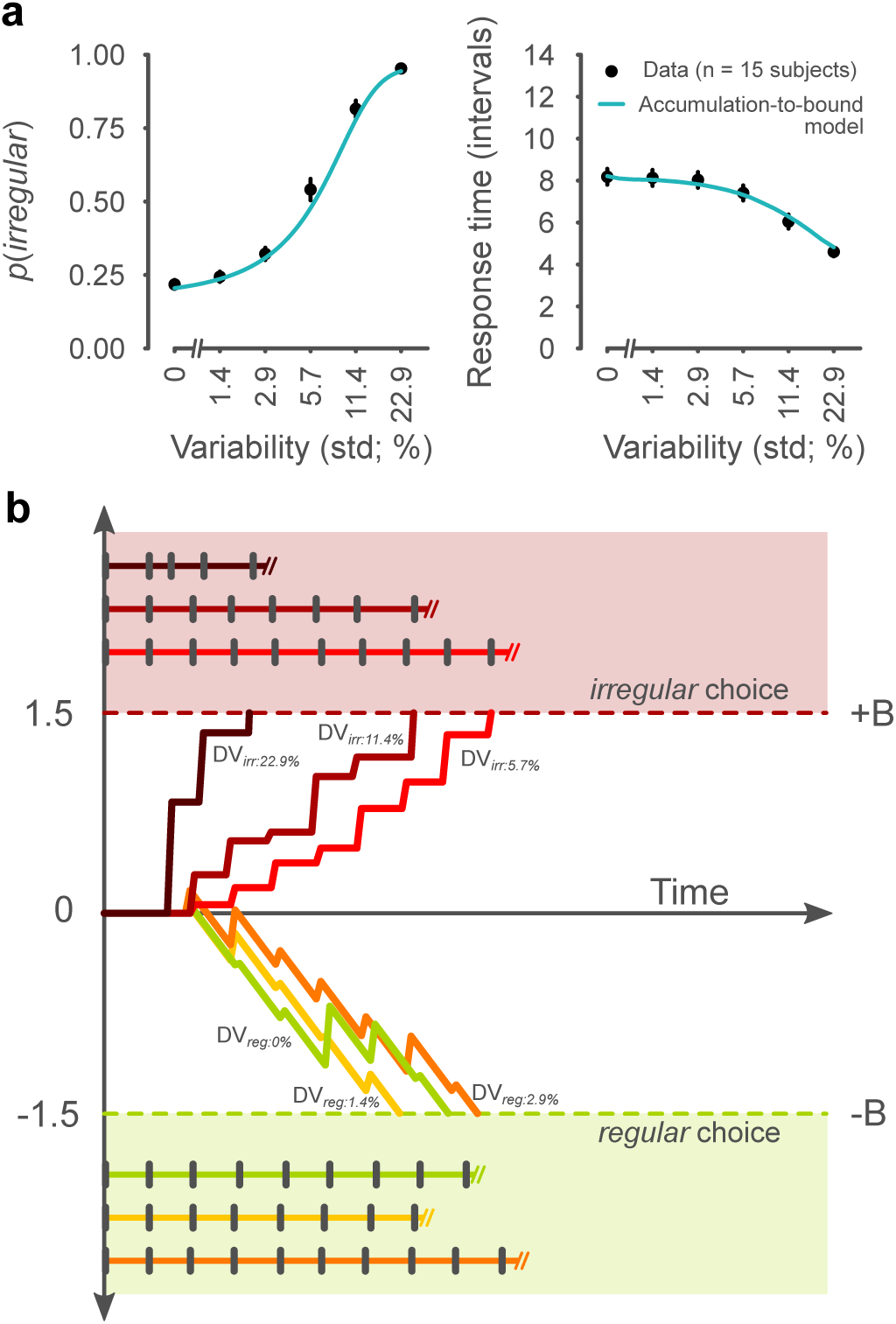
Model fit to psychometric and chronometric data. **(a)** Choice and decision times were fit with the model of decision-making (Equations 1-3). Dots and error bars illustrate across-participant mean and s.e.m. **(b)** Examples trials illustrating the model performance for irregular (*DV*_*irr*_) and regular (*DV*_*reg*_) decisions. Note how the trial with high interval variability (22.9%) leads to a fast *irregular* decision. Low variability often results in *regular* decisions (0, 1.4, 2.9%). Accumulation of irregularity evidence (*DV*_*irr*_) occurs with no leak.

**Figure 4.**
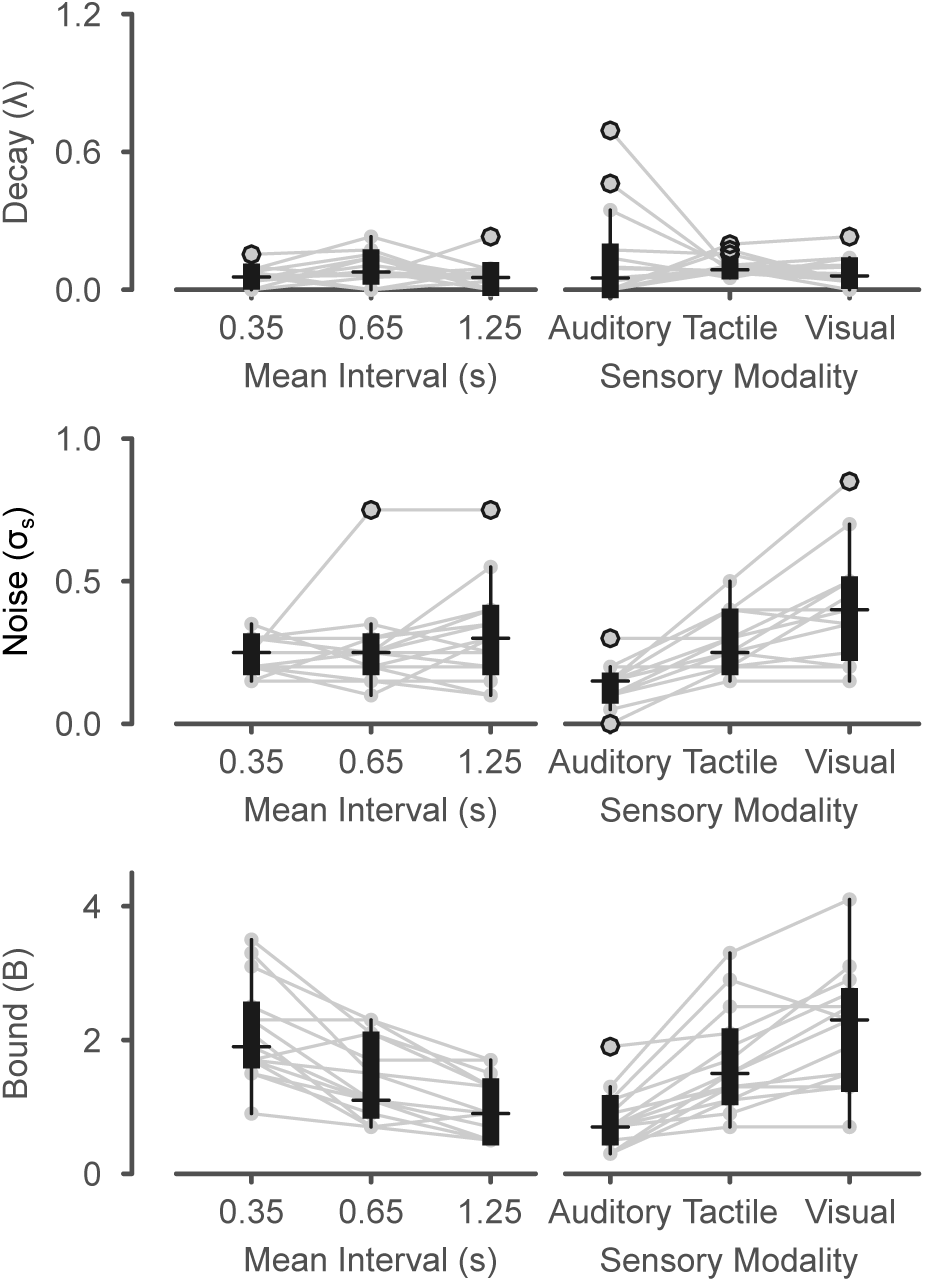
Model parameters across mean interval (0.35, 0.65, 1.25 s) and sensory modalities (auditory, tactile, visual). Leak parameters *λ* (upper panel; Equation 2), noise parameters σ_*s*_(middle panel; Equations 2-3), and bound height parameters *B* (lower panel) are plotted for each experimental condition (n=15 participants). Lines connect the same participant across conditions. Boxplots illustrate range, interquartile range, median, and outliers (open circles).

## Results

### Discrimination accuracy and response time

Subjects performed the regularity discrimination task in which they decided whether a train of sensory pulses had regular or irregular interpulse intervals (Figure 1a). Importantly, they freely communicated their choice at any time after stimulus onset. As expected, trains of stimuli with highly variable intervals were easily discriminated as *irregular*, and the decisions were made faster as variability increased (Figure 2a). The psychometric curves showed a sigmoidal increase in the proportion of *irregular* responses as a function of increasing irregularity. It is important to note that 22 ±1% (mean ±s.e.m; n=15 participants) of all regular trials (0% irregularity) were incorrectly classified as irregular by the participants (termed *false alarms* in the signal detection theory framework). This an expected result indicating that regular stimuli were often confounded with stimuli having very low temporal variability. In other words, this result reveals that participants were not certain that a train of pulses perceived as regular was in fact regular, or instead had a very small irregularity value.

Choice response times are illustrated by the chronometric curves (Figure 2b). The curves show the decreasing response times as a function of increasing interval irregularity. The chronometric curves also revealed longer response times for stimuli with longer interval durations. This tendency illustrates the fact that evidence acquisition is slower for longer interval durations.

### Discrimination accuracy scales according to the scalar property of timing

As expected from the scalar property of timing, stimuli with short interpulse intervals (0.35 s) required small deviations to be perceived as irregular, while trains with larger intervals (0.65 s, 1.25 s) required larger variability to be correctly discriminated as irregular (Figure 2a). By expressing irregularity as a percentage with respect to mean interval we observed that the psychometric curves lie close to each other, indicating that task difficulty was similar across interval lengths (Figure 2c).

Across sensory modalities, the psychometric curves showed that subjects were significantly better at discriminating auditory stimuli, as compared to tactile and visual stimuli (Figure 2e). The overall proportion of correct responses was 74 ±2%, 66 ±2% and 63 ±2% for auditory (A), tactile (T), and visual (V) stimuli, respectively (auditory performance significantly higher than tactile and visual; paired t-tests; A-T comparison: *t*(449)=8.1, p=7×10^−15^; A-V comparison: *t*(449)=8.3, p=1×10^−15^). As illustrated by the sigmoidal fits to the psychometric data, the curve for auditory stimuli had leftward shift, and a low false alarm rate. The discrimination of tactile stimuli was marginally better compared to visual stimuli, as demonstrated by the lower false alarm rate of tactile stimuli (T-V comparison: *t*(449)=2.5, p=0.012).

### Response times do not scale across interval durations

Instead of elapsed time in seconds, the chronometric curves can be expressed as the number of elapsed intervals, thus expressing response time in interval units (Figure 2d). We had expected this normalization would yield similar response times (number of intervals) across interval durations. However, we found large differences in the number of intervals that participants observed across interval durations (Figure 2d). On average, participants perceived 11.61 ±0.14 (mean ±s.e.m.) intervals before deciding that a stimulus was regular for the 0.35 s interval trials, but they waited only 4.97 ±0.07 intervals for the 1.25 s intervals. This large difference in the number of perceived intervals was an unexpected result because in our task each interval provides a quantum of information, and we had expected subjects to make use of similar amounts of information across interval durations. As we will discuss further after the model fits, this faster than expected response times explain the lower performance levels that subjects achieved for the 1.25 s psychometric curve (Figure 1c). The faster than expected response times suggest that subjects were cutting short the longer decision times that would have arisen from longer interpulse intervals.

Finally, the chronometric curves show that response times were similar across modalities for low irregularity levels (0, 1.4, 2.9%; one-way ANOVA *F*(2,942)=2.9, p=0.051) but decreased faster for the auditory stimuli as irregularity increased (Figure 1f). This indicates that participants were faster to identify the irregularity of auditory stimuli as compared to visual and tactile stimuli (5.7, 11.4, 22.9%; one-way ANOVA *F*(2,537)=6.3, p=0.002).

### Participants accumulate irregularity evidence

To determine whether behavioral choices and response times could be explained by a strategy of accumulation of timing evidence (irregularity), we fit the decision model to the mean psychometric and chronometric data. The fits showed that the model provided a satisfactory description of the behavioral data (R^2^_psycho_=0.98; R^2^_chrono_ =0.98; Figure 3a). Importantly, the model fit demonstrated that the accumulation of irregularity evidence occurs without leak (fitted *λ =* 0; Equation 2). The absence of information decay is an important result ruling out the hypothesis that subjects detect irregularity from a single large temporal deviation. Instead of waiting for a large temporal deviation to generate an *irregular* decision, our model fit indicates that participants integrated all evidence available from the train of sensory stimuli.

Six example trials illustrating the model’s behavior are shown in Figure 3b. The three trials that ended in i*rregular* decisions (*DV*_*irr*_) show that the irregularity evidence provided by each sensory pulse is accumulated with no leak, and they also illustrate how highly variable intervals result in fast decisions. The three example *regular* decisions (*DV*_*reg*_) illustrate how when the intervals are regular the downward slope (interrupted by noise jumps) determines the response time.

### Model parameters across interval duration and sensory modalities

Finally, to gain insight into differences of the decision-making process observed across sensory modalities and interval durations, we fit the model separately to each participant under each experimental condition (Methods). Consistent with the fit to overall mean data, the leak parameter was low (Figure 4; geometric mean of *λ* =0.017, 0.006-0.031 95% bootstrap confidence interval (de la Cruz & Kreft, 2019)). In other words, the lifetime of irregularity evidence was high across sensory modalities and interval durations (lifetime 1/ *λ* = 58.8 intervals; Figure 4). No significant influence of sensory modality or interval duration over the decay parameter was observed (Kruskal-Wallis *χ*^2^(5)=11.2, p=0.048). This important result further demonstrates that the accumulation of timing evidence is a robust strategy that participants employ across sensory modalities and interval durations.

The noise parameter had similar values across interval durations (*χ*^2^(2)=1.74, p=0.42), and this is consistent with the scalar property of timing stating that variability is a constant fraction (Weber’s fraction) of interval duration. Consistent with the behavioral results, the model fits show that noise in time estimation is largest in the visual domain, and lowest for auditory stimuli (Figure 4, *χ*^2^(2)=23.7, p=7×10^−6^). This result supports the idea that sensory representations of auditory stimuli have a higher temporal accuracy as compared to the visual and tactile domains.

Interestingly, we found that the bound height parameter decreased for intervals of longer durations (Figure 4, *χ*^2^(2)=17.6, p=1.5×10^−4^). This result is related to the observation that participants perceive only ∼5 long intervals before committing to a decision (Figure 2d). The decreasing bound heights suggest that participants adjusted the bound so that progressively lower amount of information was used to commit to a decision for trains of longer intervals. In other words, participants were not willing to spend too much time gathering timing information in the longer interval trials. This interesting result reveals the otherwise hidden compromise between invested time and accuracy of sensory decisions.

The comparison of bound height across sensory modalities revealed higher bounds for visual and tactile domains, and lower bounds for auditory stimuli (Figure 4, *χ*^2^(2)=20.7, p=3.4×10^−5^). The higher bounds for the tactile and visual stimuli reveal a compensatory mechanism for the noisier timing accuracy of these sensory modalities. Higher bounds allow for more information accumulation, potentially increasing decision accuracy. However, given that response times were similar across sensory modalities (Figure 2f), and that accuracy was lower for the visual and tactile domains, we can infer that participants were not willing to invest the longer accumulation times required for the tactile and visual stimuli to reach an accuracy comparable to the auditory domain.

## Discussion

Our experiments showed that decisions about the temporal regularity of a train of sensory pulses are sustained by the accumulation of the absolute time differences between regular and regular intervals. The fits of a decision-making model to behavioral data showed that the accumulation-to-bound process has no leak, so that the evidence of each interpulse interval contributes to the final decision about temporal regularity. This rules out the strategy of detecting irregularity by waiting for a large deviant interval. Our results are consistent with the hypothesis that the brain perceives a train of sensory pulses as regular when observed onset times match the predicted onset times. In this manner, differences between observed and predicted times are interpreted as evidence favoring an irregular rhythm. In our model, irregularity evidence is calculated by the difference between the duration of the current and previous interval. However, subjects may use a non-linear function of this difference to update the value of the irregular decision variable, as has been demonstrated recently in an interval reproduction task (Egger & Jazayeri, 2018). In future research, using a non-linear function of errors could help improve the fits of our model to the behavioral data.

Consistent with previous observations, we found that participants are better at perceiving the temporal structure of auditory stimuli as compared to tactile and visual stimuli (Barne et al., 2018; Di Luca & Rhodes, 2016; Mayer et al., 2014; Merchant et al., 2015; Murai & Yotsumoto, 2016; Rammsayer, 2014; Rammsayer et al., 2015; Westheimer, 1999). This suggests differences in the temporal fidelity of the sensory representations of train of pulses across modalities. Our model captured this difference with an increased noise parameter for visual stimuli as compared to tactile and auditory stimuli. We know the visual system excels at the spatial analysis of scenes, so a lower timing precision was expected. It is important to note, however, that the model fits demonstrate that accumulation of timing evidence is a decision-making strategy shared by the auditory, tactile, and visual domains.

The behavioral results showed that the response times do not scale across the three interval lengths (0.35, 0.65, 1.25 s) as we had expected. Given that each sensory pulse delivers a quantum of timing information (they define the duration of intervals) we had expected that response time, measured in number of elapsed intervals, would be similar across interval durations. Although longer intervals resulted in longer response times (Figure 2b), this did not translate into an equal number of perceived intervals across durations. This indicates that subjects were not willing to wait too long to gather the same evidence for the long intervals as they did for short intervals (Figure 2d). This translated into a lower level of correct responses for the 1.25 intervals as compared to the 0.35 and 0.65-s intervals. Quick decision times at the expense of choice accuracy reveals a decision-making policy in which participants tried to invest similar decision times across interval durations. Closely related to this, we also found that response times are very similar across sensory modalities (Figure 2f). Given the noisier time estimates of the visual domain, subjects would have needed to increase the decision time (i.e. gather more evidence) to compensate. Our experiments did not aim to study the time-accuracy tradeoff inherent to decision-making (Hanks et al., 2014; Palmer et al., 2005; Reddi & Carpenter, 2000), but the tendency to homogenize response time across experimental conditions is a decision policy that is worth of further study.

## Acknowledgements

We thank Mehrdad Jazayeri, Hugo Merchant and Pavel Rueda for useful discussions and comments on this work. We thank Edgar Bolaños for technical assistance; Jessica González Norris for proofreading the manuscript; Luis Aguilar, Alejandro de León, Carlos Flores, and Jair García of the National Laboratory of Advanced Scientific Visualization (LAVIS). This research was supported by Dirección del Personal Académico de la Universidad Nacional Autónoma de México (PAPIIT IN207818) and CONACYT (254313, 247200). ME was funded by the Mexican Council of Science and Technology (CONACYT 453852). ME is a doctoral student from Programa de Doctorado en Ciencias Biomédicas, Universidad Nacional Autónoma de México (UNAM). VdL received support from MIT International Science and Technology Initiatives.

